# A toolset of constitutive promoters for metabolic engineering of *Rhodosporidium toruloides*

**DOI:** 10.1101/592774

**Authors:** Luísa Czamanski Nora, Maren Wehrs, Joonhoon Kim, Jan-Fang Cheng, Angela Tarver, Blake A. Simmons, Jon Magnuson, Miranda Harmon-Smith, Rafael Silva-Rocha, John M. Gladden, Aindrila Mukhopadhyay, Jeffrey M. Skerker, James Kirby

## Abstract

**Background:** *Rhodosporidium toruloides* is a promising host for the production of bioproducts from lignocellulosic biomass. A key prerequisite for efficient pathway engineering is the availability of robust genetic tools and resources. However, there is a lack of characterized promoters to drive expression of heterologous genes for strain engineering in *R. toruloides*.

**Results:** Our data describes a set of native *R. toruloides* promoters, characterized over time in four different media commonly used for cultivation of this yeast. The promoter sequences were selected using transcriptional analysis and several of them were found to drive expression bidirectionally. We measured promoter expression strength by flow cytometry using a dual fluorescent reporter system. From these analyses, we found a total of 20 constitutive promoters (12 monodirectional and 8 bidirectional) of potential value for genetic engineering of *R. toruloides*.

**Conclusions:** We present a list of robust and constitutive, native promoters to facilitate genetic engineering of *R. toruloides.* This set of thoroughly characterized promoters significantly expands the range of engineering tools available for this yeast and can be applied in future metabolic engineering studies.

## 1. Background

With the aim of replacing many petrochemical-derived fuels and chemicals with renewable alternatives, the field of metabolic engineering is moving towards the development of microbial hosts that use lignocellulosic biomass as feedstocks for bioproduction. The oleaginous, carotenogenic yeast *Rhodosporidium toruloides* (also known as *Rhodotorula toruloides*) is a promising host for sustainable bioproduction due to its innate capacity to efficiently utilize both hexoses and pentoses present in lignocellulosic biomass, as well as its capacity to grow on lignin derived intermediates and aromatics, such as *p*-coumaric acid and ferulic acid [1]. So far, much of the work on metabolic engineering or bioproduct formation in *R. toruloides* has focused on lipid accumulation [2,3]. However, the high lipid content indicates a suitability for applications to any acetyl-CoA-derived product, from fatty alcohols and methyl ketones to terpenes, with many applications such as fuels, chemicals, and pharmaceuticals [1,4–6].

To improve prospects for more advanced metabolic engineering in this fungal host, the development of robust genetic tools and resources is an immediate priority [7]. A major need on the tools front is for efficient and stable promoters to express genes of interest and selection markers. A growing branch of synthetic biology involves the pursuit and improvement of standard parts that are reliable, orthogonal and robust for non-conventional hosts [8,9]. A small selection of promoters has already been characterized to modulate expression of heterologous genes in *R. toruloides*, the majority of them being metabolite-responsive promoters rather than constitutive [7,10–13]. While promoters that are responsive to certain metabolites are valuable tools, a toolset of well characterized constitutive promoters remains necessary to explore the full potential of strain engineering. In this context, functional bidirectional promoters would be a valuable addition to the toolset, since they can be convenient for optimization of multi-gene pathways.

The most common method used for transformation of *R. toruloides* is *Agrobacterium tumefaciens*-mediated transformation (ATMT), a useful strategy for genomic integration of heterologous genes but is a laborious and time-consuming process [14,15]. For this work, we adapted a high-efficiency chemical transformation method from the one developed for *Saccharomyces cerevisiae* [16] to streamline the engineering workflow for *R. toruloides*. We performed RNA sequencing analysis to select promoters from *R. toruloides* that are likely to result in high- and medium-level constitutive expression. Several of these promoters were predicted to be bidirectional. We characterized these promoters by cloning them in a dual-fluorescent reporter system and monitored expression in four different media over a cultivation period of seven days. Of the set of promoters tested, several appear to be good candidates for driving heterologous expression in this host, thus expanding the toolset available for genome editing.

## 2. Results

### 2.1. Construction and characterization of promoter library

RNA-sequencing data derived from *R. toruloides* cultivated on several different carbon sources was used to identify constitutive promoters driving high and medium expression levels of native genes (predicted expression levels are classified as described in the Methods section). All selected promoters are considered to be constitutive, based on variance in transcript abundance across six different growth conditions. Of the 29 promoter regions selected, 13 comprise a complete intergenic sequence (i.e., between start codons of two neighboring divergent genes) and were therefore identified as putative bidirectional promoters. Each promoter was cloned into a reporter gene construct consisting of mRUBY2 and EGFP in two orientations, designated as orientation 1 and 2 (Figure 1A and Figure SI1). A strain containing the reporter gene cassette lacking a promoter was used as the negative control, referred to here as ∆*car2*. In total, 59 constructs were transformed and integrated into the *R. toruloides* genome, 29 constructs in orientation 1 and 29 in orientation 2 (Figure 1A), and one control with no promoter.

**Figure 1.**
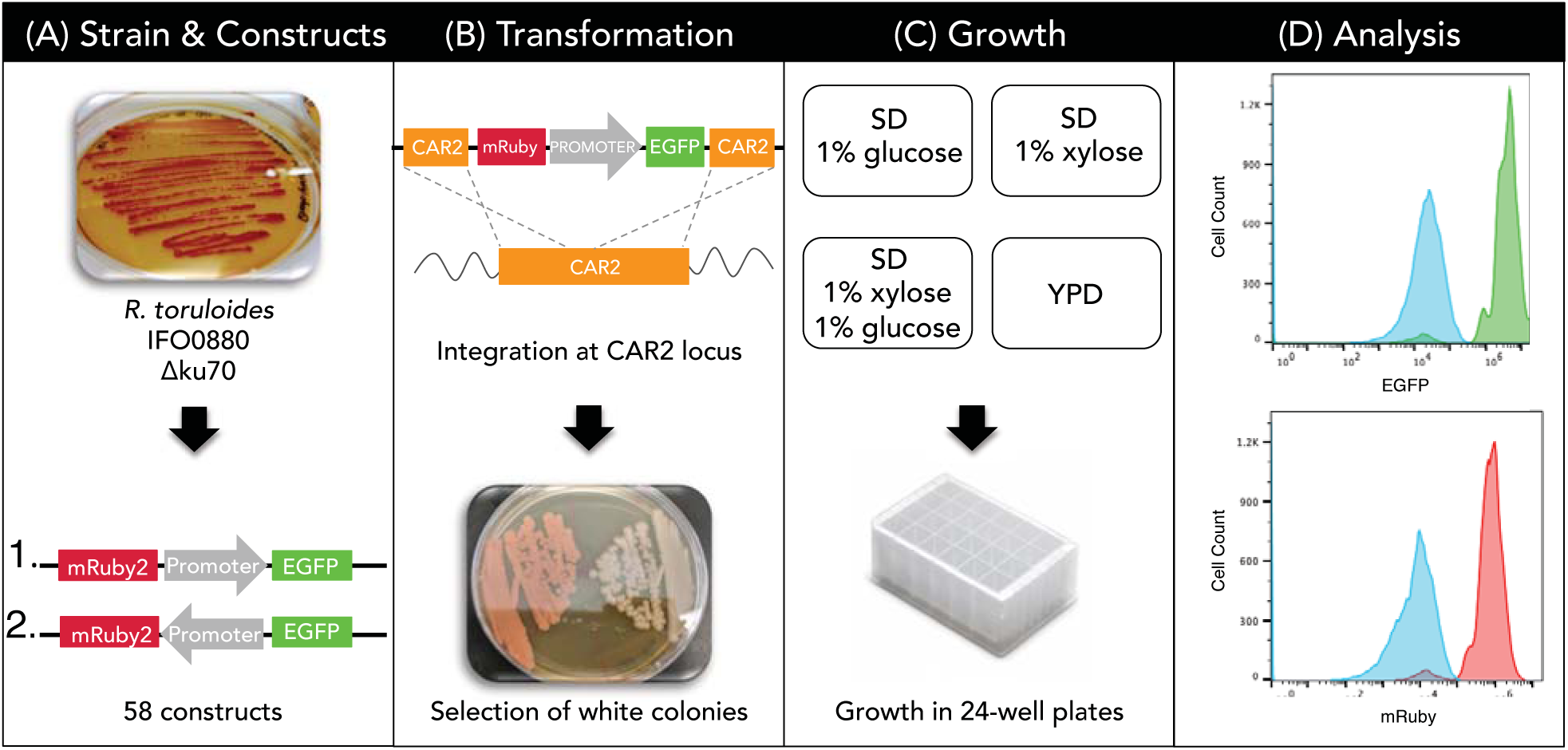
Experimental workflow. **(A)** A *KU70*-deficient strain of haploid *R. toruloides* IFO0880 was used for all experiments. Each promoter-reporter cassette was constructed in two orientations (1 and 2, above) in order to construct strains that report expression of both EGFP and mRuby2 from every promoter. 13 of the 29 promoters were predicted to be bidirectional (not shown). **(B)** Constructs were integrated into the *CAR2* locus of the *R. toruloides* genome using a lithium-acetate-based transformation method. Strains harboring a correctly-targeted construct were determined by a white Δ*car2* phenotype and fluorescence. **(C)** The Δ*ku70* parent strain, a control ∆*car2* strain, and three transformants for each of the 58 promoter strains were cultivated in 24-deep-well plates in four different media: YPD, SD supplemented with 1% (w/v) glucose, SD supplemented with 1% (w/v) xylose, and SD supplemented with 1% (w/v) each of glucose and xylose. Cultures were grown for 7 days, with sample collection at 8, 24, 48, 96 and 168 hours. **(D)** Reporter gene expression was analyzed by flow cytometry. The upper and lower histograms show fluorescence from EGFP (upper, green histogram) and mRuby2 (lower, red histogram) driven by promoter P9, compared to the control ∆*car2* strain (blue histograms); these populations were sampled at 24 hours in SD supplemented with 1% (w/v) each of glucose and xylose.

The workflow from construction of expression constructs to promoter analysis by flow cytometry is shown in Figure 1. Each construct was flanked by 1 kb sequences homologous to the *R. toruloides CAR2* gene, which encodes a bifunctional phytoene/lycopene synthase [10,11]. Disruption of the *CAR2* gene through homologous recombination results in white colonies due to lack of carotenoid production (Figure 1B). Three confirmed transformants for each construct were inoculated into 24-deep-well plates containing LB medium, grown overnight, and then sub-cultured into the various test media at a starting OD_600_ of 0.05. One rich medium (YPD) and three defined media (SD supplemented with 1% glucose, 1% xylose, or 1% of each sugar) were selected to represent typical media compositions [1] to culture *R. toruloides* (Figure 1C).

To select appropriate time points for reporter fluorescence measurements, we monitored optical density at 600 nm (OD_600_) and measured extracellular sugars by high performance liquid chromatography (HPLC) in all four media. Representative data, for the parental strain, *R. toruloides* IFO0880 ∆*ku70* and a strain harboring reporter construct P14, can be found in Figure SI4 and SI5, respectively. Samples were collected at 8, 24, 48, 96 and 168 hours for measurement of reporter fluorescence by flow cytometry. For constructs containing bidirectional promoters, fluorescence of both reporters was measured simultaneously.

### 2.2. Monodirectional promoters

Of the 16 putative monodirectional promoters selected for investigation, 12 resulted in medium to high fluorescence when driving EGFP expression in *R. toruloides*. Hierarchical clustering of a heatmap generated from flow cytometry data (Figure 2) shows that the strongest monodirectional promoters under most conditions are P14 (translation elongation factor 1, TEF1), P17 (hypothetical protein, RTO4_15825), and P15 (mitochondrial adenine nucleotide translocator, ANT) – see Table 1 for more information on promoter sequences. Overall, promoters P14, P17, P15, P4, P27, P18, and P10 are classified as strong promoters, while promoters P7, P1, P5, P3 and P6 are clustered as medium-strong promoters (in decreasing order of strength). It is worth noting that promoter P7, which natively drives expression of GAPDH and is commonly viewed as a benchmark strong promoter, is classified as medium-strong within this promoter set.

**Table 1.**
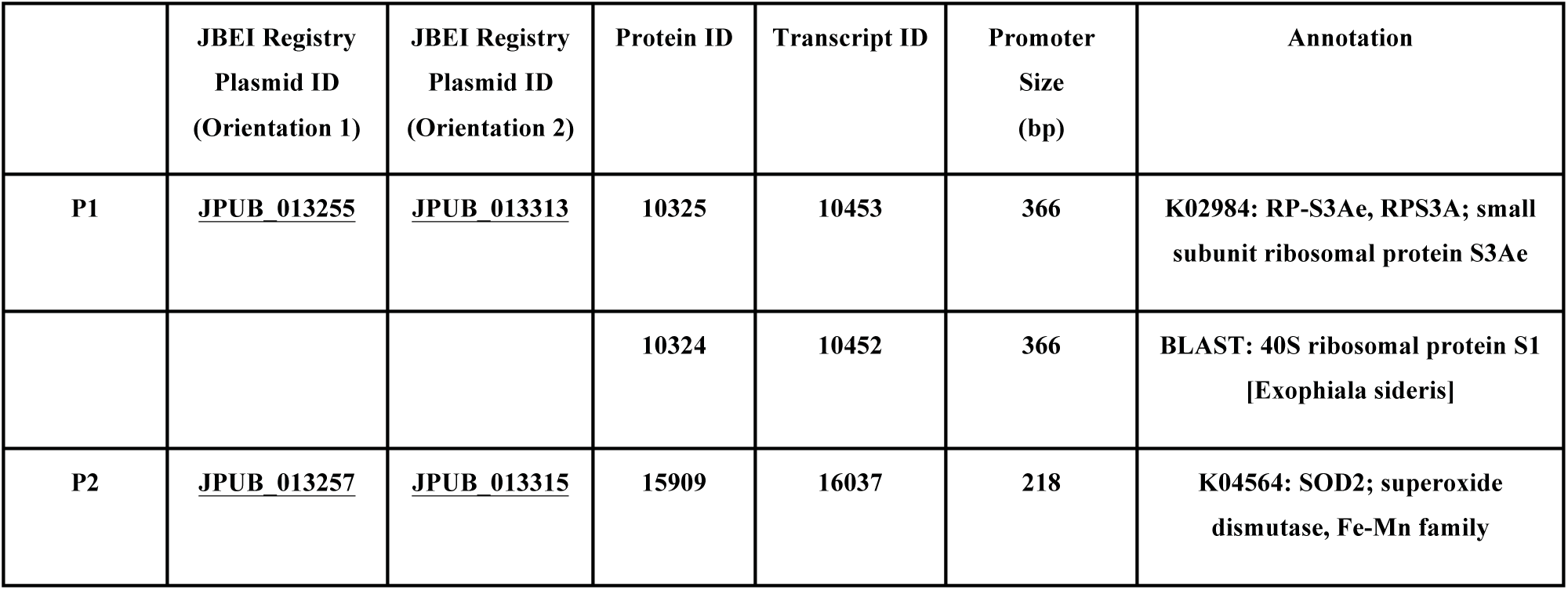
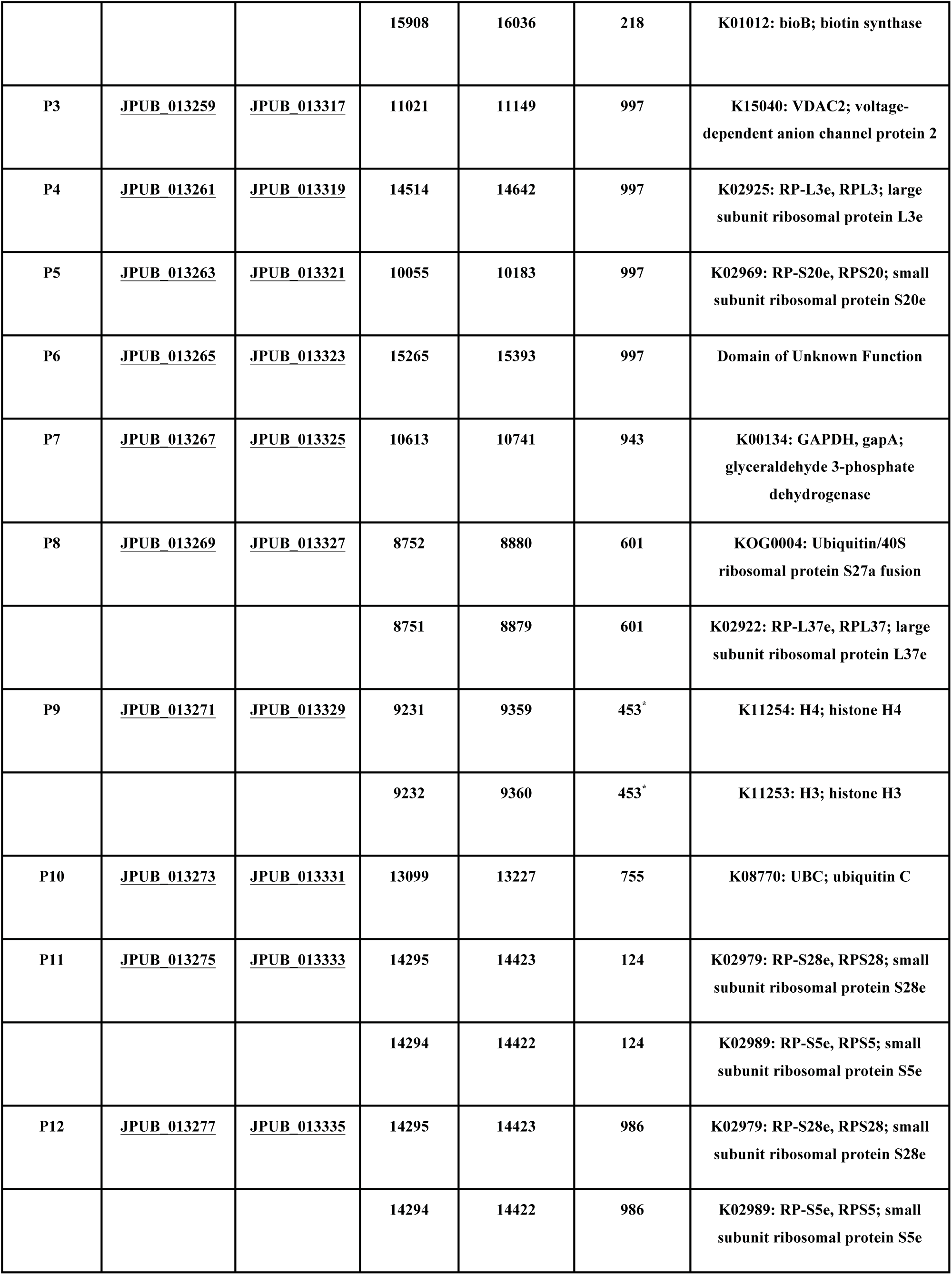
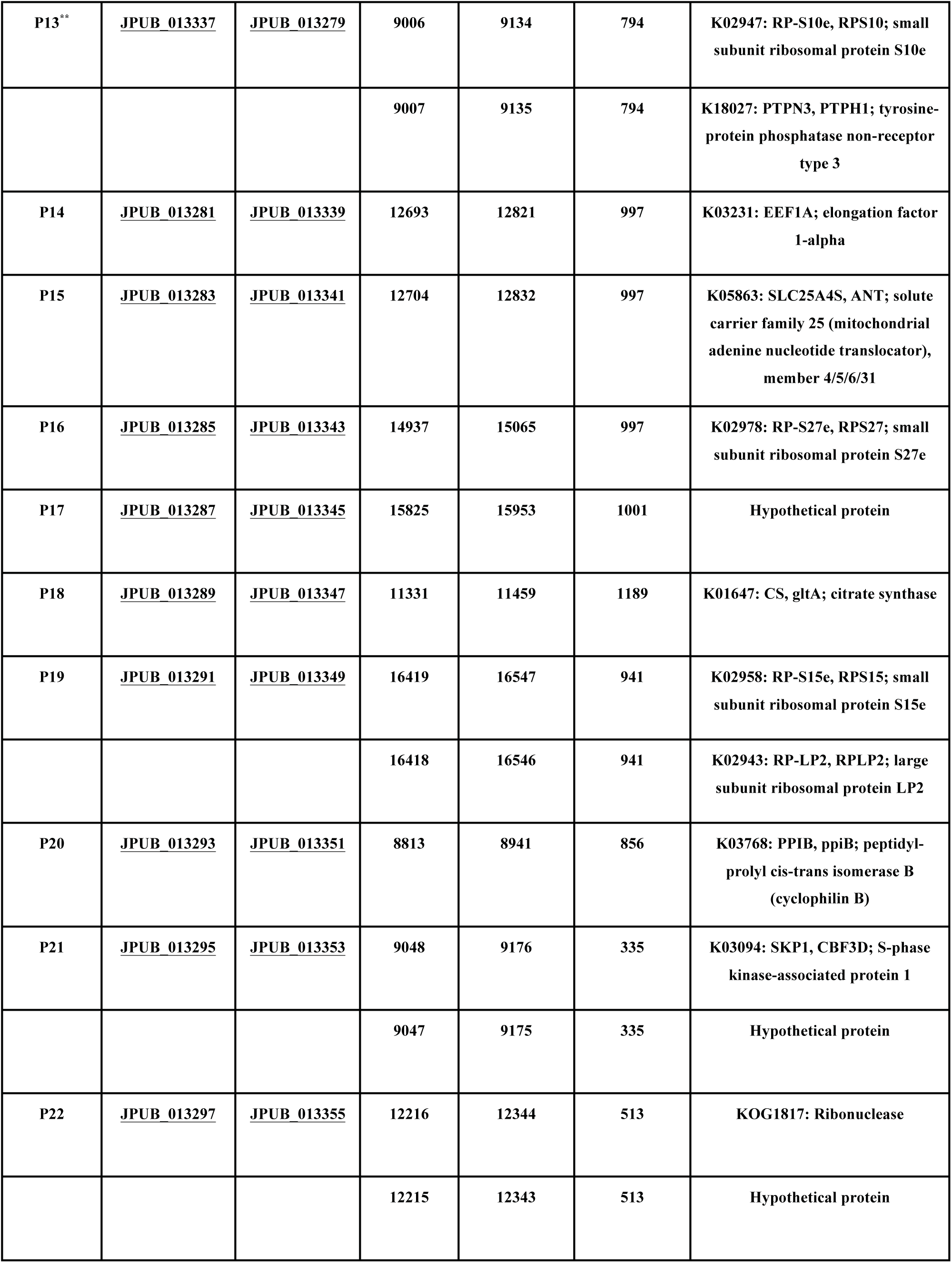
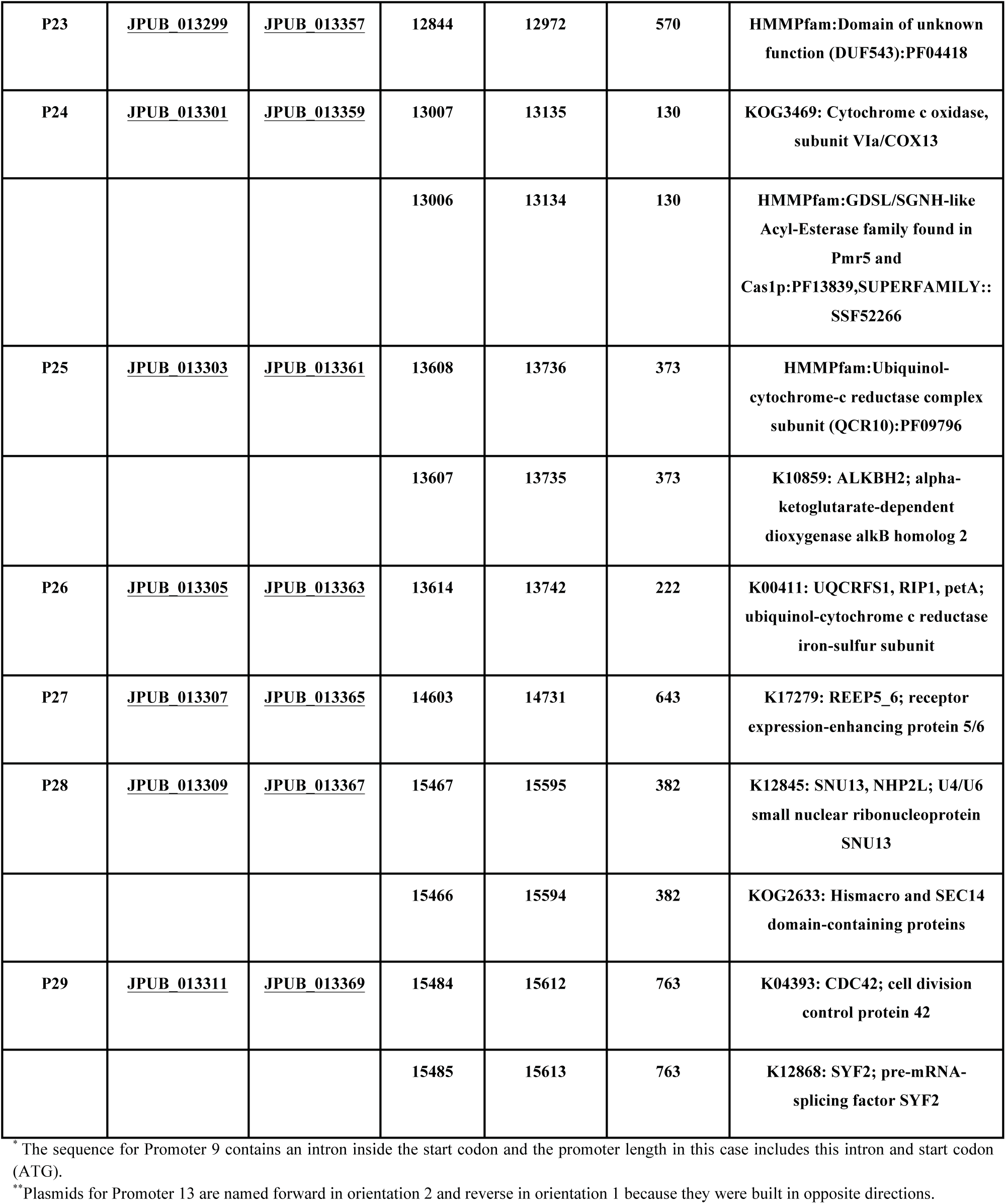
Promoters investigated and strains generated in this study. Promoters selected from RNA-sequencing data are listed with their locations in the *R. toruloides* genome, links to their respective constructs in the Joint BioEnergy Institute (JBEI) registry, and annotations of putative gene functions. Promoter pairs that span complete intergenic regions and are therefore predicted to be bidirectional are represented by two listings, annotated as P1 and P1R, etc. Protein ID and Transcript ID refers to the *Rhodosporidium toruloides* IFO0880 v4.0 genome sequence, which can be found on the Joint Genome Institute (JGI) MycoCosm site: https://genome.jgi.doe.gov/Rhoto_IFO0880_4/Rhoto_IFO0880_4.home.html.

**Figure 2.**
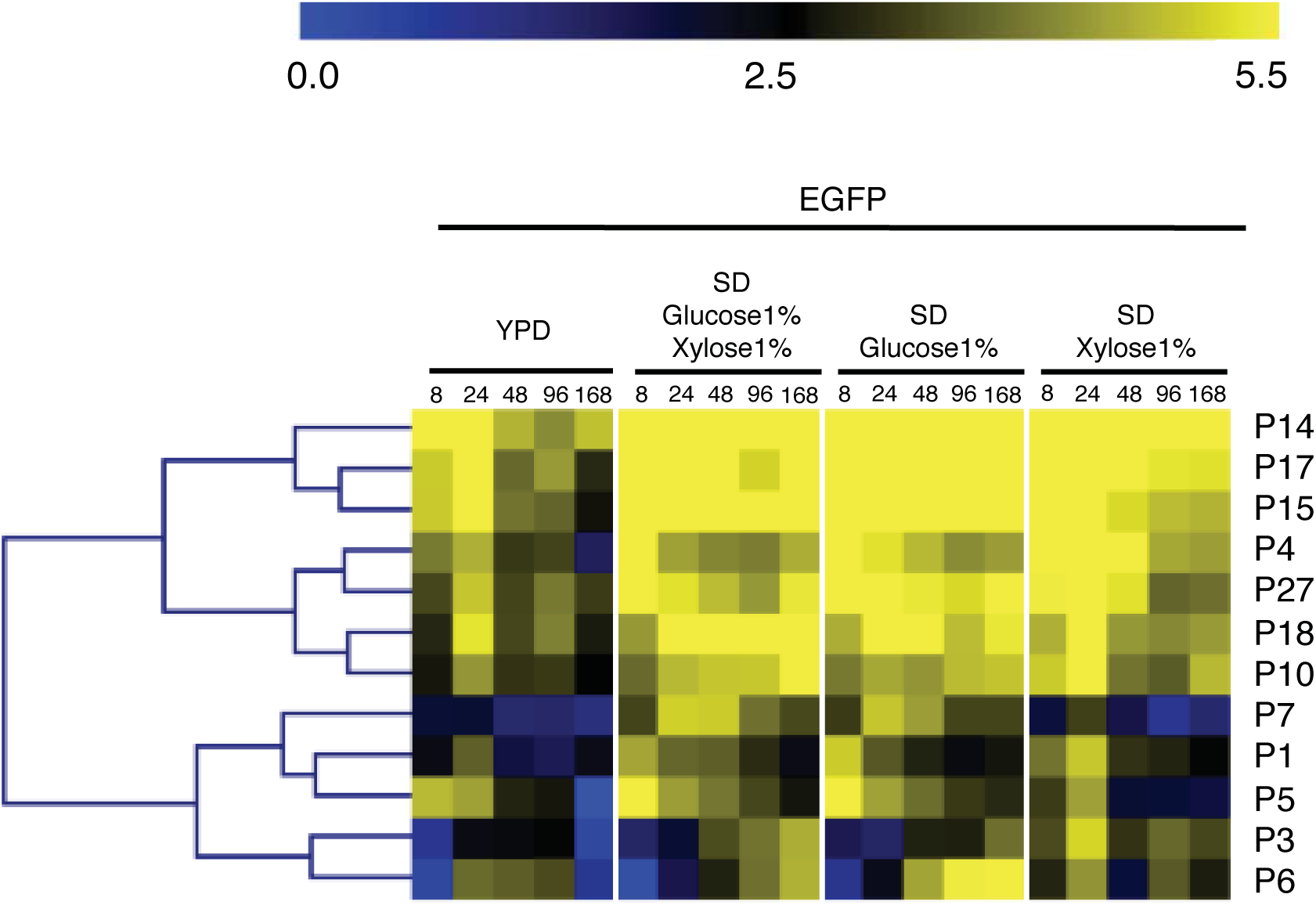
Expression of EGFP from monodirectional promoters. Fluorescence was measured by flow cytometry of cell populations sampled at five time points from four different media for strains harboring promoter constructs in orientation 1. The heatmap was generated using MultiExperiment Viewer (MeV), and promoters were clustered by hierarchical clustering using Euclidean distance. Fluorescence expression values are on a Log2 scale (color scale is shown in the upper bar), calculated as described in Methods. Each column represents a time point, shown in hours. Promoters represent each row with their names on the right side of the heatmap.

In order to evaluate consistency of expression of different coding sequences, promoters were also used to drive expression of mRuby2 (Figure SI2). Again, promoters P6, P3 and P1 were classified as medium-strong promoters and promoters P14, P15 and P17 were clustered as the strongest promoters.

### 2.3. Bidirectional promoters

Each of the 13 promoters that was predicted to be bidirectional (i.e., selected by transcriptomics and comprising a complete intergenic sequence in *R. toruloides*) was positioned in between the divergent coding sequences for EGFP and mRuby2, allowing expression in both directions to be monitored simultaneously. Of these, 8 resulted in medium-to-high reporter fluorescence in both directions in at least one condition. Hierarchical clustering of a heatmap generated from flow cytometry data shows that the strongest bidirectional promoter pairs are P9 and P9R (histones H3 and H4, respectively), P12 and P12R (small subunit ribosomal proteins S28e and S5e, respectively), and P19 and P19R (ribosomal proteins S15e and LP2, respectively) (Figure 3). The forward (EGFP) and reverse (mRuby2) promoter strengths for these three bidirectional pairs are well-balanced across a range of conditions. Under most conditions, promoters P9 and P9R, constituting the strongest bidirectional promoter pair, are similar in strength to the strongest of the monodirectional promoters. Promoter pairs P13/R, P28/R, P22/R, P29/R and P8/R were clustered as medium-strength promoters in at least one direction. Of these, P29 and P29R are the most evenly-balanced, while P8 and P8R are the most divergent in terms of promoter strength. P8R (ribosomal protein, L37e) is significantly stronger than P8 (Ubiquitin/40S ribosomal protein S27a fusion), across most of the conditions tested.

**Figure 3.**
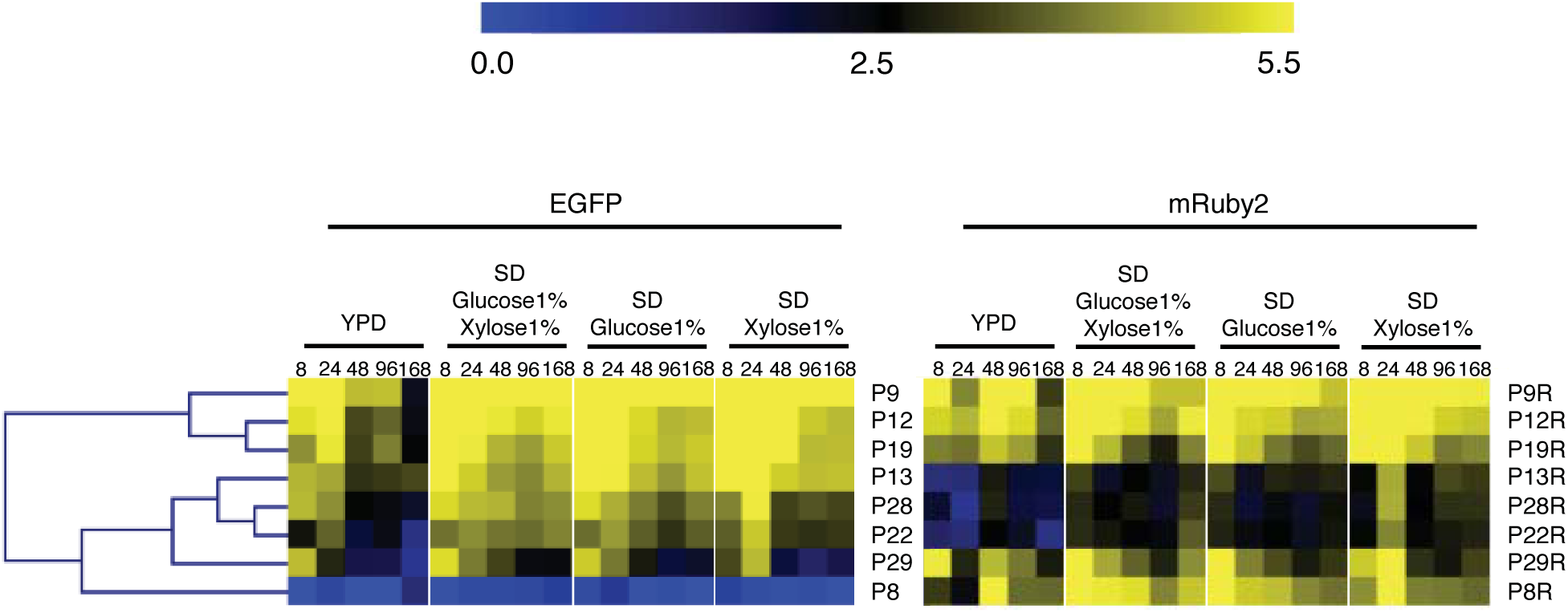
Expression of EGFP and mRuby2 from bidirectional promoters. Fluorescence of EGFP and mRuby2 for each bidirectional promoter pair was measured in strains harboring promoter constructs in orientation 1. The heatmap was generated using MultiExperiment Viewer (MeV), and promoters were clustered by hierarchical clustering using Euclidean distance. Fluorescence expression values are on a Log2 scale (color scale is shown in the upper bar), calculated as described in Methods. Each column represents a time point, shown in hours. Promoters are represented in rows with their names on the right side of the heatmap. All time points for all media are shown for both EGFP and mRuby.

Fluorescence from a second reporter was measured for each of these bidirectional promoters by cloning them in orientation 2 (Figures SI1 and SI3). Promoters P9/R, P19/R and P12/R were again classified as the strongest bidirectional promoter pairs while P8/R and P29/R were overall the weakest promoter pairs. Promoter P22 was classified as medium bidirectional promoter when assayed in both orientations.

### 2.4. Correlation of mRuby2 and EGFP expression

To evaluate promoter reliability and robustness, the expression level of the two reporter genes (EGFP and mRuby2) driven by the same promoter were compared. Linear regression and correlation analysis of fluorescence from the two reporters was performed for all the promoters comparing EGFP from orientation 1 and mRuby2 from orientation 2 (EGFP expression for these promoters is shown in Figures 2 and 3 and mRuby2 expression is shown in Figures SI2 and SI3) in the four different media at 48 hours (Figure 4).

**Figure 4.**
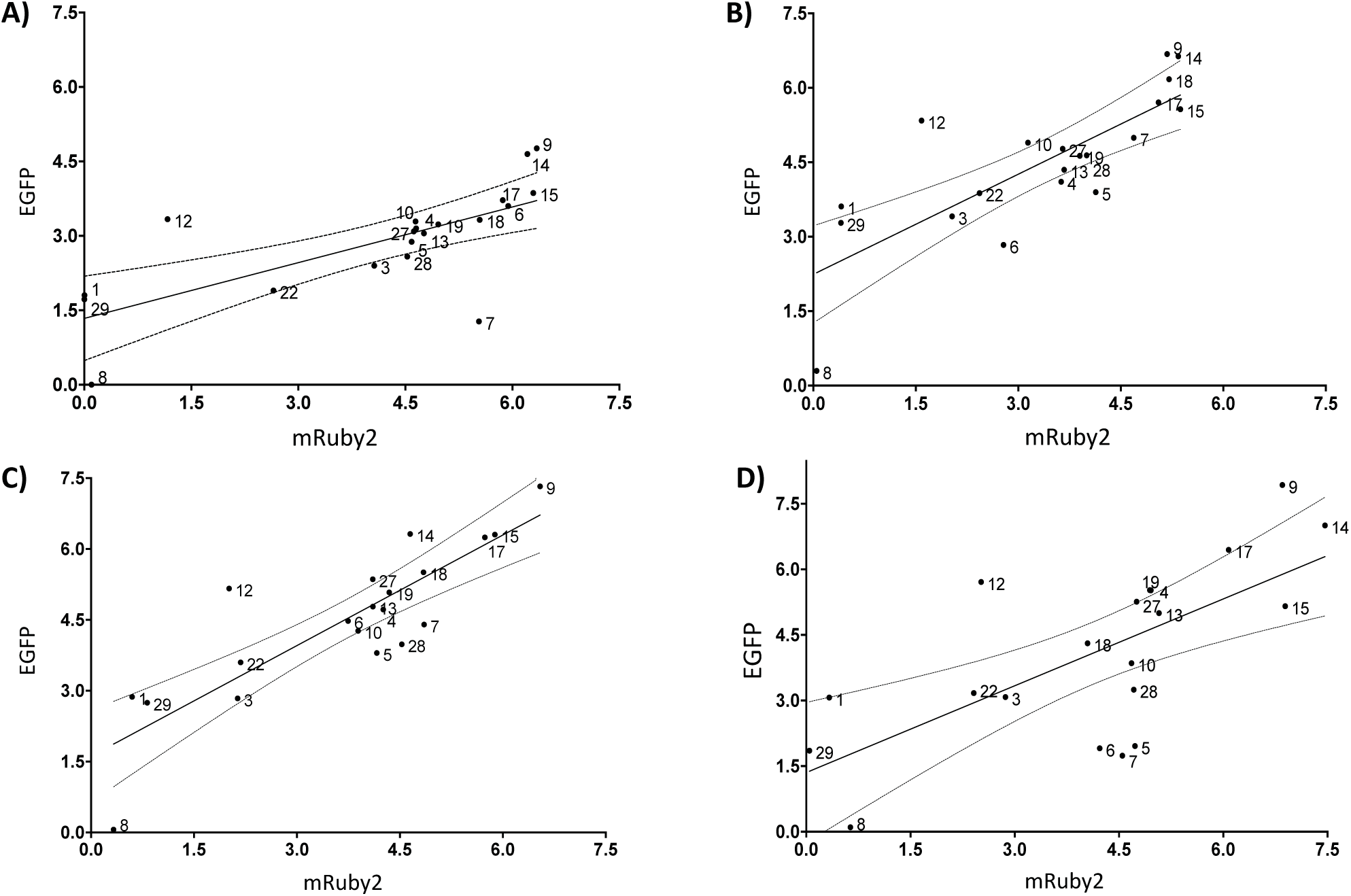
Correlation of fluorescence from two reporters expressed from *R. toruloides* promoters. To get a sense of promoter robustness, each promoter was cloned in front of two reporter genes, EGFP and mRuby2 (denoted as orientation 1 and 2 in Figure 1). Average fluorescence output for both reporters at 48 hours is charted for each promoter in four different media: correlation between EGFP expression from strains containing constructs in orientation 1 and mRuby2 expression from strains containing constructs in orientation 2. Solid line is the linear regression and dashed lines are the 95% confidence interval. YPD (A); SD supplemented with 1% (w/v) each of glucose and xylose (B); SD supplemented with 1% (w/v) glucose (C); and SD supplemented with 1% (w/v) xylose (D). Pearson correlation for promoter expression in SD with 1% glucose and 1% xylose resulted in a R^2^ of 0.6159 and for SD 1% glucose resulted in a R^2^ of 0.7223, both with a P value of < 0.0001. Correlation of promoter expression in SD 1% xylose resulted in a R^2^ of 0.471 and in YPD resulted in a R^2^ of 0.5041, both with a P value of 0.0005.

Promoter P1, P12, and P29 expressed EGFP at significantly higher levels than mRuby2. This could suggest a composability effect, in which contextual effects such as the mRNA secondary structure surrounding the ATG is less favorable for EGFP than for mRuby2 [17–19]. The benchmark promoter P7 (GAPDH) expressed mRuby2 at higher levels than EGFP in strains grown in YPD medium or SD with 1% xylose, indicating that expression levels from this promoter may be less predictable in certain media. Most of the strongest promoters, including P4, P9, P10, P14, P15, P17, P18 and P27, produce similar levels of EGFP and mRuby2 fluorescence, and are recommended for metabolic engineering needs.

## 3. Discussion

The lack of standard parts that can be reliably used for genetic manipulation is often a bottleneck in metabolic engineering studies involving non-model microbial hosts. We aimed to find and characterize a collection of promoters to support reliable gene expression in the fungal host *R. toruloides*. We selected 29 *R. toruloides* promoters, based on transcriptomics studies, and examined them in four different media over seven days using a dual-reporter system.

P_*GAPDH*_ is one of the most widely used promoters for engineering a variety of yeasts, including *R. toruloides*, where it was employed for expression of heterologous terpene synthases [1]. However, 13 of the promoters investigated in this study were found to be stronger than P_*GAPDH*_, under the majority of conditions (Figures 2 and 3).

Temporal changes in promoter strength can be an important factor in guiding their selection for pathway engineering. Under control of several of the promoters, reporter fluorescence reaches its maximum early (at 8 or 24 hours) and then diminishes over time or drops to a lower but stable level (Figures 2 and 3). Interestingly, this occurs for two of the strongest promoters, P15 (*ANT*) and P9 (histone H3) when strains are cultured in YPD medium, but not in the SD media. This is likely due to the cells reaching stationary phase significantly faster in YPD compared to SD media (Figure SI4 and SI5). In contrast, expression from promoters P3 (voltage-dependent anion channel protein 2, *VDAC2*) and P6 (unknown function) increases over time under some conditions. These two promoters may be useful for applications where late expression of a protein is desired (e.g., generation of metabolic products that are inhibitory to growth or consumed during log phase).

Correlation of fluorescence from EGFP and mRuby2 at the 48 hours suggests that most of the promoters express both reporter genes at similar rates, suggesting a reasonable level of composability, with a few exceptions (Figure 4). A similar analysis of fluorescence from the two reporters driven by promoter P9, which includes every time point and growth medium, indicates that the expression level of the two reporters is similar under all conditions for this promoter (Figure SI6).

We investigated promoters from *R. toruloides* that were posited to have medium or strong transcriptional activity in both directions. Bidirectional promoters can reduce design and assembly complexity and also help keep DNA construct sizes within reasonable limits, a useful feature when engineering multigene pathways [20–22]. Of the 14 promoters that were predicted to be bidirectional according to RNA-sequencing data (P1, P2, P8, P9, P11, P12, P13, P19, P21, P22, P24, P25, P28 and P29) fluorescence of both reporters was detected for 8 of them, indicating bidirectional transcription (Figure 3). Expression in the forward and reverse directions from the strongest bidirectional promoter pairs (P9/P9R, P12/P12R and P19/P19R) is balanced (Figures 3 and SI3), a feature that should prove valuable for metabolic engineering applications where equal expression of two genes is sought. Each of these three promoters drives expression of two genes that are closely related in function. Promoters P9 and P9R, the bidirectional pair natively responsible for expression of histones 3 and 4, respectively (Table 1), was shown to be the strongest promoter of the entire collection, under most conditions.

Promoters P6, P9, P10, P12, P14 and P15 contain introns in their predicted 5ʹ UTRs, a feature often associated with increased promoter strength. Introns can increase transcript levels by affecting the rate of transcription, nuclear export, transcript stability and even mRNA translation [23]. In *S. cerevisiae* it was shown that intronic sequences had a positive impact on expression of ribosomal proteins, both transcriptionally and post-transcriptionally [24].

So far, there have been 3 reports of relatively constitutive promoters for use in *R. toruloides*, totaling a set of 8 promoters: *GAPDH* (also called *GPD1*), *FBA1*, *PGK1*, *PGL1*, *TPI1*, *LDP1*, *ACC1* and *FAS1* [10,11,13,14]. Since normalization of the reporter genes used to characterize these promoters differs between these studies, it is challenging to compare strength of these promoters with the collection presented here.

Synthetic biology studies often rely heavily on the fine-tuning of gene expression levels, to achieve a metabolic balance and a high product titer, rate and yield [5,9,25,26]. The majority of engineering work to date performed to date in *R. toruloides* has relied on random-integration strategies such as ATMT [1,2,14,15], that offer little control over integration locus or copy number. The strong promoters characterized in this work will complement the recent development of CRISPR technology for *R. toruloides*, allowing for site-specific integration while maintaining high expression rates [27].

## 4. Conclusions

RNA sequencing data was found to be a useful starting point for identification of *R. toruloides* promoters that can be used for heterologous expression. The collection of 12 monodirectional and 8 bidirectional native promoters presented in this work is the largest promoter set published for *R. toruloides*, including the first bidirectional promoters reported for this organism. Among these were 13 promoters that are stronger than the benchmark GAPDH promoter, the most commonly used promoter for yeasts. This characterized promoter set expands the *R. toruloides* genetic toolbox and is likely to be valuable for future metabolic engineering efforts in this promising host.

## 5. Methods

### 5.1. Media and growth conditions

Synthetic defined (SD) medium was made using Difco yeast nitrogen base (YNB) with ammonium sulfate and without amino acids (Becton, Dickinson & Co., Sparks, MD) and complete supplemental mixture (CSM; Sunrise Science Products, San Diego, CA), following manufacturer’s directions. Lysogeny broth (LB) was made using Difco LB (Miller) mixture (Becton, Dickinson & Co.) and yeast peptone dextrose (YPD) medium was made as normal, including 2% peptone, 1% yeast extract (Becton, Dickinson & Co.) and 2% glucose. Xylose and glucose were from Sigma-Aldrich (St. Louis, MO).

For the promoter studies, all 60 *R. toruloides* IFO0880 *∆ku70* strains (the parent strain, 58 strains containing promoter constructs, a *∆car2* control strain) were inoculated into 24-deep-well plates containing 2 mL of LB (Miller) and grown overnight at 30 °C with shaking at 200 rpm. These cultures were then used to inoculate (at a 1:100 dilution) four different media in 24-deep-well plates: SD 1% xylose, SD 1% glucose, SD 1% xylose plus 1% glucose, and YPD. These cultures were grown to exponential phase overnight at 30 °C with shaking at 200 rpm and were then used to inoculate 3 mL of identical media in 24-deep-well plates at a starting OD_600_ of 0.05. These cultures were grown for 7 days at 30 °C in a shaker at 200 rpm with samples taken every 8, 24, 48, 96 and 168 hours.

### 5.2. Strains and plasmid construction

The parent strain for all work described here is *R. toruloides* IFO0880 harboring a deletion for the non-homologous end joining (NHEJ) gene, *KU70*, to facilitate a higher proportion of site-specific integration events in each transformation (Figure 1A) [2]. *R. toruloides* strain IFO0880 *∆ku70*, a haploid strain, was used for transformations to favor homologous recombination. Strains and plasmids used in this study are available upon request through the Joint BioEnergy Institute Strain Registry (https://public-registry.jbei.org/). Gene synthesis and plasmid construction were performed by Joint Genome Institute (JGI) and two constructs were designed for each promoter so that it was positioned between EGFP and mRuby2 coding sequences in both directions. Constructs also contain a Nourseothricin acetyltransferase (*nat1*) cassette, conferring resistance to Nourseothricin, and homology regions corresponding to the *CAR2* locus. A schematic representation of the expression cassette built for integration into *R. toruloides* genome can be seen in Figure 1 and selected promoters and their combined annotations can be seen in Table 1.

### 5.3. RNA-sequencing

RNA-sequencing was performed at the Joint Genome Institute and used to identify a set of promoters that drive high or medium expression in different media including SD medium with 2% glucose (NCBI accession SRP183741, SRP183740, SRP183739), and M9 minimal medium with added trace elements and 2% glucose (NCBI accession SRP164139, SRP164140, SRP164141) or 10 mM p-coumaric acid (NCBI accession SRP164142, SRP164143, SRP164144). Another dataset was generated at the University of California, Berkeley and included in the analysis (NCBI accession GSE128360), which consists of samples grown in YNB minimal medium with 3% glucose and different carbon to nitrogen ratios (7mM ammonium chloride or 168mM ammonium chloride), and a library made of pooled samples grown in several media including YNB medium with different carbon sources (glucose, xylose, glycerol, or glutamate) and YPD medium with different conditions (growth phase, osmolarity, temperature, and oxygen availability). Sequenced reads were trimmed and filtered using BBTools (https://jgi.doe.gov/data-and-tools/bbtools/), mapped to IFO0880 genome sequence using HISAT2 [28], mapped reads were assigned to genes using featureCounts [29], and read counts were used to calculate Fragments Per Kilobase of transcript per Million mapped reads (FPKM). High expression was defined as log2 of the mean FPKM values above 10, and medium expression was defined as log2 of the mean FPKM values between 8 and 10. In order to select promoters with constitutive expression, 20 genes with the lowest variance across different growth conditions were selected from each high and medium expression gene. Subsequently, promoter regions of high and medium expression genes were selected for testing, spanning 1 kb of upstream sequence or a shorter distance in cases where an upstream CDS was identified. In some cases, the predicted promoter region of a selected gene was cut short by an upstream CDS on the opposite strand and was annotated as a putative bidirectional promoter.

### 5.4. *R. toruloides* transformation

Plasmids were purified from *E. coli* DH10B strains using a QIAprep plasmid miniprep kit (Qiagen, Redwood City, CA, USA) and digested with *Pvu*II (FastDigest, Thermo Scientific, Waltham, MD, USA) to release each construct from the vector backbone (with exception of constructs 5, 14, 18, 34, 43 and 47 that were amplified by PCR using Phusion High Fidelity DNA Polymerase (Thermo Scientific, Waltham, MD, USA) since they had *Pvu*II sites inside their promoter sequences). The transformation method was adapted from Gietz [16]. *R. toruloides* IFO0880 *∆ku70* strain was inoculated into 10 mL of YPD and incubated overnight at 30 °C with shaking at 200 rpm. Then, the culture was diluted to an OD_600_ of 0.2 in fresh YPD and incubated at 30 °C on a shaker at 200 rpm until it reached an OD_600_ of 0.8. The cultures were harvested, centrifuged at 3000 × g for 10 min, washed once with 5 mL of water and then resuspended in 1 mL 100 mM lithium acetate (pH 7.5) (LiAc) in 1.7 mL tubes. The cells were centrifuged again and the LiAc was removed. Next, the transformation mixture was added, consisting of 240 µL PEG 4000 (50% w/v), 36 µL 1.0 M LiAc (pH 7.5), 10 µL single-stranded salmon sperm DNA (10 mg/mL) and 5 µg of the transforming DNA in 74 µL of water. After mixing by vortexing, cells were incubated at 30 °C for 30 min. Subsequently, 34 µL DMSO was added and cells were heat shocked at 42 °C for 15 minutes. Cells were then centrifuged at 7,200 rpm in a microfuge and the supernatant was removed by pipetting. Cells were resuspended in 1.5 mL YPD, transferred to culture tubes, and allowed to recover overnight at 30 °C at 200 rpm. The cells were plated in YPD agar plates containing 100µg/mL of Nourseothricin Sulfate (G-Biosciences, St. Louis, MO) and incubated at 30 °C for two days. The plates were then stored at 4 °C for two days to differentiate the white from orange colonies. For each construct, at least three independent transformants were grown overnight in YPD and tested for fluorescence by flow cytometry (Figure 1B).

### 5.5. Flow cytometry

High-throughput flow cytometry experiments were performed using the Accuri C6 flow cytometer equipped with an autosampler (BD Accuri™ RUO Special Order System, Becton, Dickinson and Company, New Jersey, USA). All fluorescence measurements were performed on biological triplicates. A total of 30,000 events were recorded at a flow rate of 35 µL/min and a core size of 16 µm for each sample. Both fluorescent proteins were excited at 488 nm, EGFP emission was detected at 530 nm and mRuby2 emission was detected at 675 nm. Data acquisition was performed as described in the Accuri C6 Sampler User’s Guide. The acquired data was analyzed in the FlowJo^®^ software (Becton, Dickinson & Company), where populations were gated and the median of all fluorescence curves were obtained.

### 5.6. Statistics

For the fluorescence values, the median value of each fluorescence curve of each replicate was extracted using FlowJo^®^. The mean of the triplicates was calculated from the median values, as well as the standard deviation (STD) and the coefficient of variation (CV). The CVs of all promoters were normalized to a scale of 0 to 1. The resulting means were then divided by the mean fluorescence of the negative control ∆*car2* to normalize all samples in relation to background fluorescence on the EGFP and mRuby2 channels. The resulting normalized values were then transformed to log_2_ values represented in the figures as Relative Fluorescence Units (RFU). Heatmaps were assembled using Multiple Experiment Viewer (MeV), where promoters were clustered by hierarchical clustering using Euclidean distance as the distance metric selection through average linkage clustering. The remaining graphs were created using GraphPad Prism^®^ (GraphPad Software, San Diego, CA), also used to calculate linear regression as needed.

### 5.7. Growth rate experiments

For growth rate experiments, strains were grown in the same conditions as the fluorescence measurement experiments. Samples were taken at 0, 8, 24, 48, 96 and 168 hours after inoculation. OD_600_ was monitored using a SpectraMax Plus 384 spectrophotometer (Molecular Devices, San Jose, CA, USA).

### 5.8. High performance liquid chromatography (HPLC) analysis

Sugars were quantified on an Agilent Technologies 1200 series HPLC (Agilent Technologies, Santa Clara, CA, USA) equipped with an Aminex HPX-87H column (BioRad, Hercules, CA) as described previously [30]. Samples were filtered through 0.45 µm filters (VWR, Visalia, CA, USA) before injection of 5 µl of each sample onto the column. Sugars were monitored by a refractive index detector, and concentrations were calculated by integration of peak areas and comparison to standard curves for the compounds of interest.

## 6. Declarations

### 6.1. Ethics approval and consent to participate

Not applicable.

### 6.2. Consent for publication

Not applicable.

### 6.3. Availability of data and material

Strains generated and analyzed here are available in The Joint BioEnergy Institute Inventory of Composable Elements (JBEI-ICE) at https://public-registry.jbei.org/folders/437.

### 6.6. Authors’ contributions

JMG and JMS and conceived the project. MW, James Kirby, LCN, and AM designed the experiments. LCN performed the experiments, data analysis and wrote the manuscript. Joonhoon Kim performed the RNA sequencing analysis. JFC, AT, and MHS constructed all plasmids. James Kirby, JMS, Joonhoon Kim, MW, AM, RSR, JMG, and BAS critically revised the manuscript. All authors read and approved the final manuscript.

## 6.7. Acknowledgements

The authors would like to thank Samuel Coradetti for sharing RNA sequencing data, Di Liu and Gina Geiselman for the helpful comments and Sophie Comyn for assistance with confocal microscopy of *R. toruloides*. Strains and plasmids are available in The Joint BioEnergy Institute Inventory of Composable Elements (JBEI-ICE) at https://public-registry.jbei.org/folders/437.

## 10. SUPPLEMENTARY INFORMATION

**Figure SI1.**
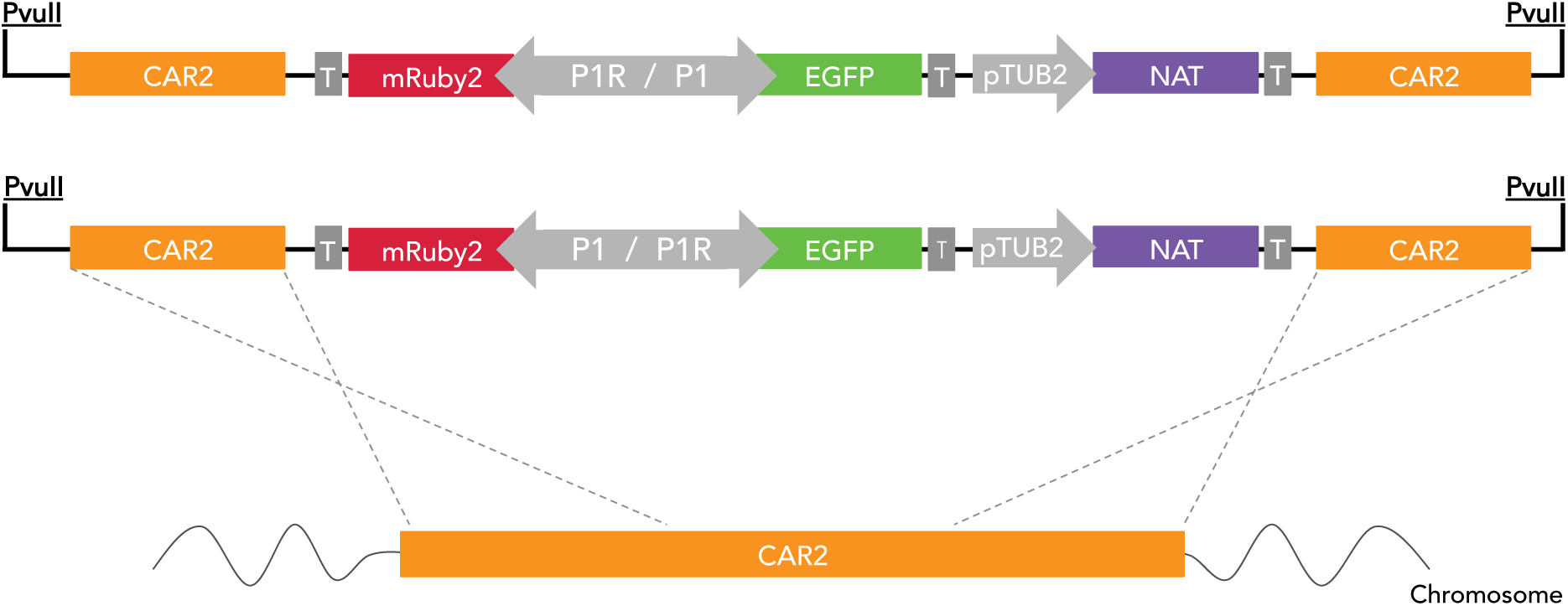
Schematic representation of the expression cassettes to be inserted into the *R. toruloides* genome through homologous recombination. Representative design of the constructs in orientation 1 (top), showing promoter P1 driving EGFP expression and P1R driving mRuby2 and orientation 2 (lower), where the promoter fragment has been reversed. Promoter-reporter constructs, paired with a *NAT* cassette (conferring resistance to nourseothricin), are flanked by *CAR2* gene fragments to facilitate integration of the cassette into the *CAR2* locus of the *R. toruloides* chromosome.

**Figure SI2.**
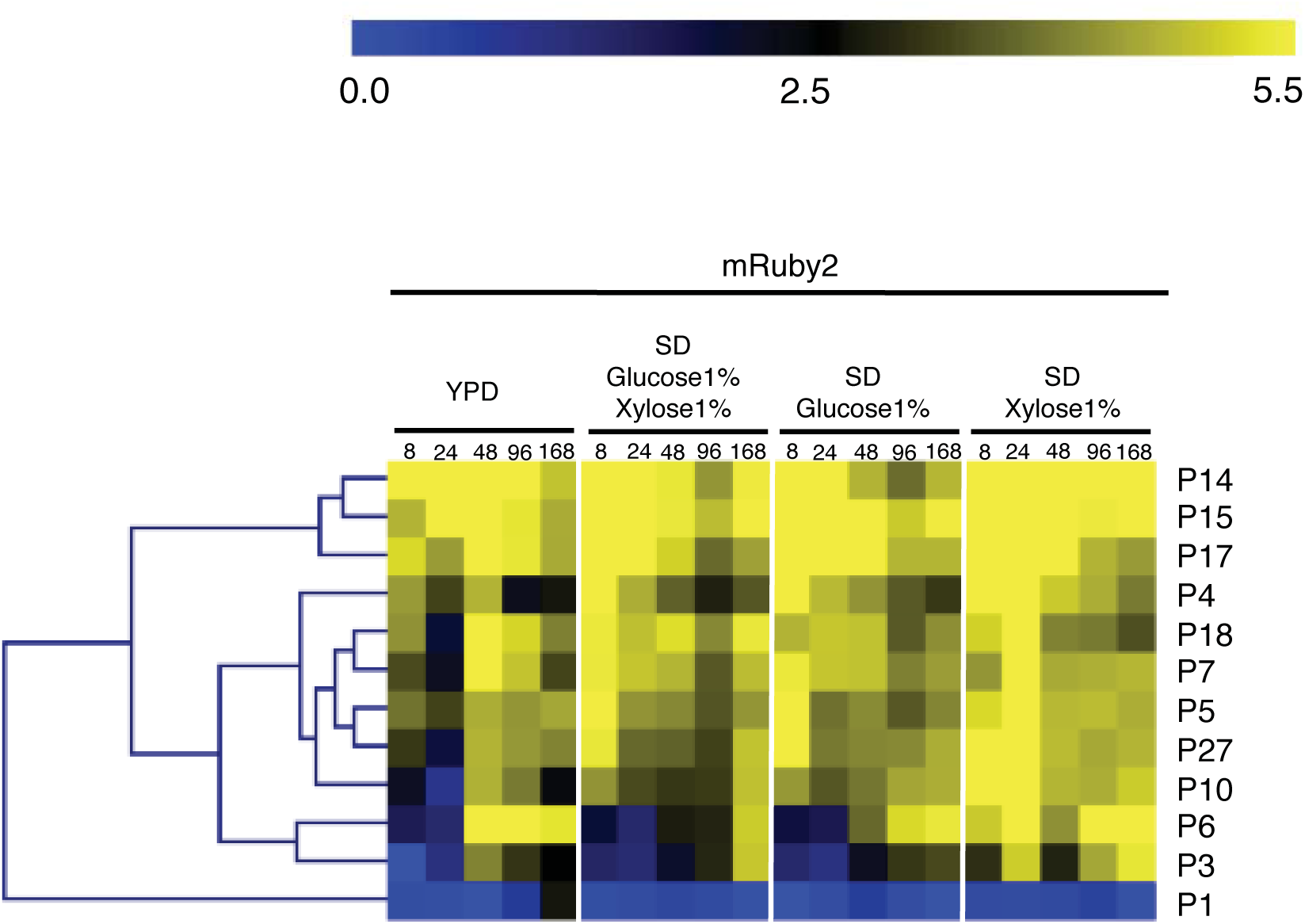
Expression of mRuby2 from monodirectional promoters in orientation 2. Fluorescence was measured by flow cytometry of cell populations sampled at five time points from four different media. The heatmap was generated using MultiExperiment Viewer (MeV), and promoters were clustered by hierarchical clustering using Euclidean distance. Fluorescence expression values are on a Log2 scale (color scale is shown in the upper bar), calculated as described in Methods. Each column represents a time point, shown in hours. Promoters are represented in rows with their names on the right side of the heatmap.

**Figure SI3.**
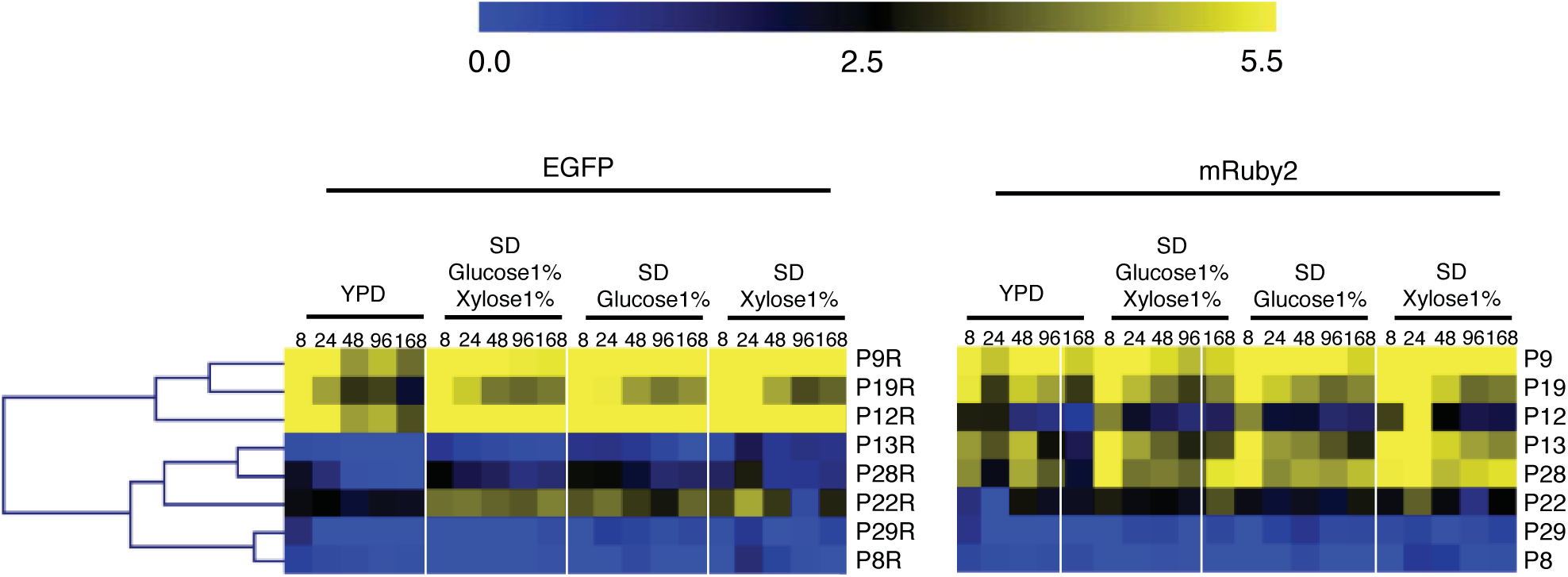
Expression of EGFP and mRuby2 from bidirectional promoters, in orientation 2. Fluorescence of EGFP and mRuby2 for each bidirectional promoter pair in orientation 2 was measured by flow cytometry of cell populations sampled at five time points from four different media. The heatmap was generated using MultiExperiment Viewer (MeV), and promoters were clustered by hierarchical clustering using Euclidean distance. Fluorescence expression values are on a Log2 scale (color scale is shown in the upper bar), calculated as described in Methods. Each column represents a time point, shown in hours. Promoters are represented in rows with their names on the right side of the heatmap.

**Figure SI4.**
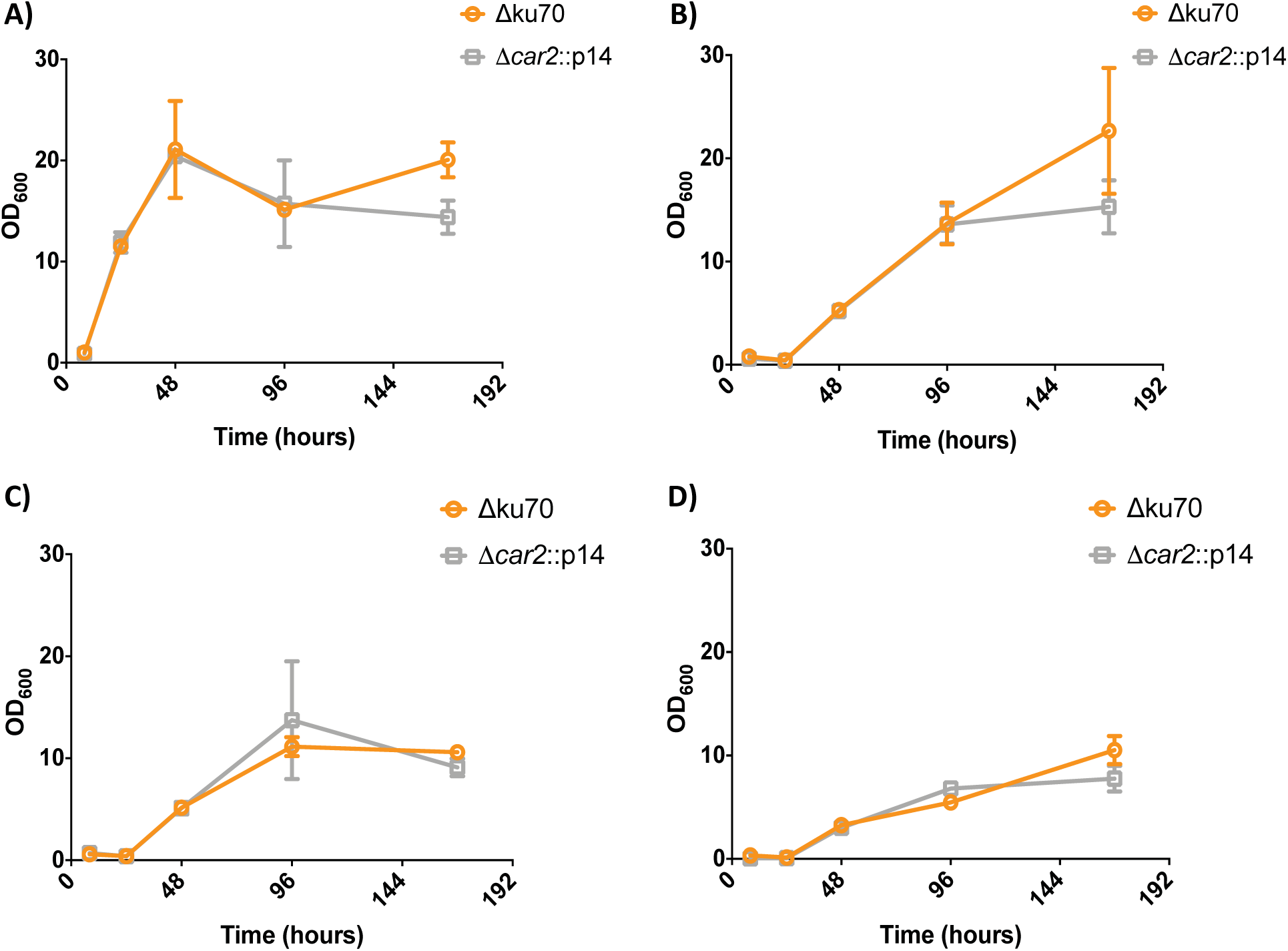
Growth of the parental *R. toruloides* ∆*ku70* strain and a strain harboring promoter construct P14. Growth was monitored by OD_600_ measurements, taken at 8, 24, 48, 96 and 168 hours from strains grown in: YPD (A); SD supplemented with 1% (w/v) each of glucose and xylose (B); SD supplemented with 1% (w/v) glucose (C); and SD supplemented with 1% (w/v) xylose (D).

**Figure SI5.**
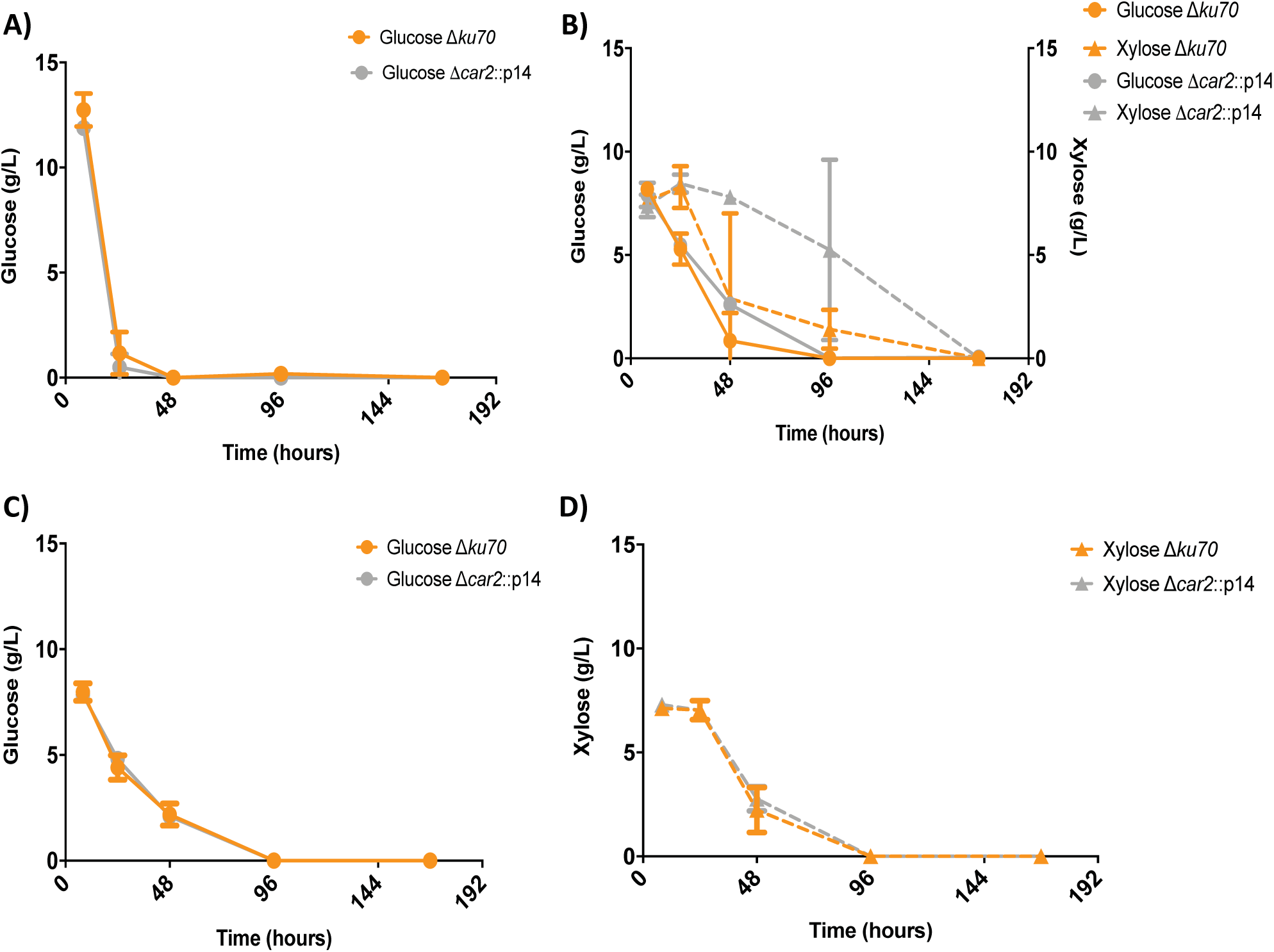
Glucose and xylose consumption in the four media used for this study. Two strains are shown: the parental *R. toruloides* ∆*ku70* strain and strain harboring promoter construct P14. Sugars were quantified by HPLC from samples taken at 8, 24, 48, 96 and 168 hours from strains grown in: YPD (A); SD supplemented with 1% (w/v) each of glucose and xylose (B); SD supplemented with 1% (w/v) glucose (C); and SD supplemented with 1% (w/v) xylose (D).

**Figure SI6.**
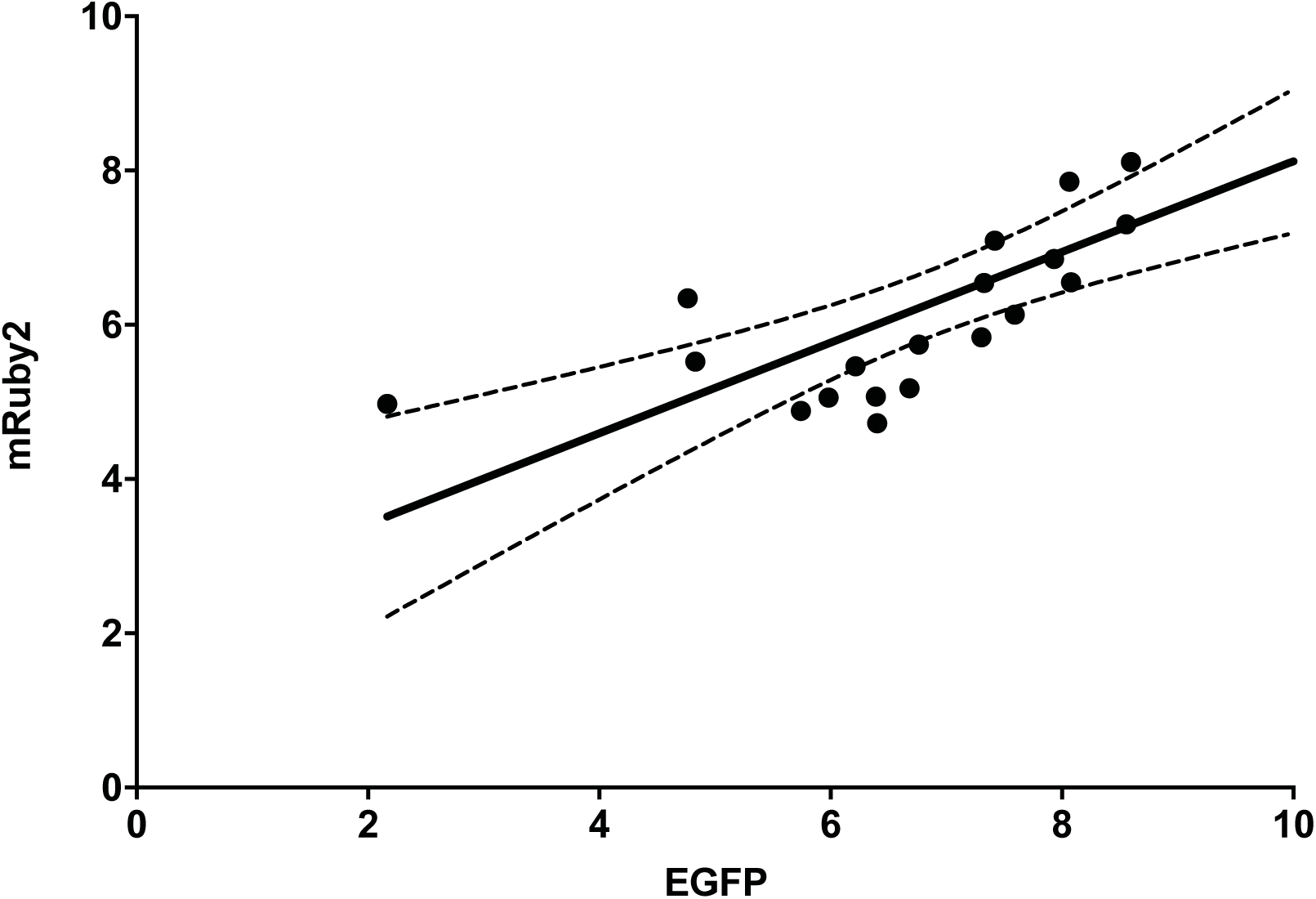
Correlation between EGFP and mRuby2 expression from promoter P9 in all 4 media and all 5 time points. Fluorescence expression values are on a Log2 scale. R^2^ = 0.5537 for a P value of 0.0002. Solid line is the linear regression and dashed lines are 95% confidence interval.

